# Hierarchical and context-dependent GR-MR signalling governs endogenous corticosteroid decoding in the heart

**DOI:** 10.64898/2026.09.09.750321

**Authors:** F. Sacchi, S. Boriati, R. Caliandro, S. Da Pra, I. Del Bono, A. Husetić, C. Bongiovanni, C. Miano, N. Pianca, S. Mazzone, A. Cabrini, R. Tassinari, C. Ventura, M. Lauriola, M. M. Gladka, G. D’Uva

## Abstract

Endogenous corticosteroids bind both the glucocorticoid receptor (GR) and mineralocorticoid receptor (MR), yet how this shared ligand input is decoded in the heart remains unclear. Previous work established that corticosterone suppresses postnatal cardiomyocyte proliferation through GR. Here we show that corticosteroid responses are governed by a functional GR-MR hierarchy and by cellular context. Genetic or pharmacological GR inhibition redirects corticosterone towards cardiomyocyte proliferation, and MR antagonism or silencing abolishes this effect. GR disruption also enhances aldosterone-induced proliferation in cardiomyocyte-enriched cultures, indicating that GR restrains MR output beyond ligand allocation. However, aldosterone fails to increase cardiomyocyte proliferation in mixed cultures and instead stimulates fibroblast proliferation and activation. By contrast, corticosterone combined with GR inhibition promotes cardiomyocyte proliferation without inducing stromal proliferation or profibrotic activation. Both treatments induce MR nuclear localisation in fibroblasts, showing that their divergent stromal effects arise despite comparable receptor nuclear engagement. Following myocardial infarction, circulating corticosterone increases and GR inhibition enhances cardiomyocyte MR nuclear localisation. In adult murine myocardium, corticosterone plus GR antagonism increases cardiomyocyte cell-cycle activity in an MR-dependent manner, with analogous responses observed in porcine and human myocardium. These findings identify hierarchical and context-dependent GR-MR signalling as a mechanism of corticosteroid decoding and a potential route to cardiac regeneration.

## INTRODUCTION

Corticosteroids are adrenal steroid hormones that comprise two major classes: glucocorticoids, which are key regulators of metabolism, stress responses, inflammation and tissue maturation, and mineralocorticoids, which mainly control electrolyte balance, blood pressure and cardiovascular homeostasis (Oakley and Cidlowski 2015; Richardson et al. 2016; Cruz-Topete et al. 2020; Galow et al. 2023). Although these hormones are often interpreted through their cognate receptors, their biological effects are shaped by overlapping receptor binding.

Physiological glucocorticoids bind not only the glucocorticoid receptor (GR), but also the closely related mineralocorticoid receptor (MR), for which they display high affinity (Hultman et al. 2005; Mifsud and Reul 2016). In classical aldosterone-responsive epithelial tissues, MR selectivity is conferred by 11β-hydroxysteroid dehydrogenase type 2 (11β-HSD2), which converts active glucocorticoids into inactive metabolites and thereby protects MR from glucocorticoid occupancy. By contrast, cardiac cells express little or no 11β-HSD2, and circulating glucocorticoids are present at concentrations more than 100-fold higher than aldosterone, suggesting that cardiac MR is likely to be predominantly occupied by endogenous glucocorticoids under physiological conditions (Richardson et al. 2016). Thus, the cardiac response to corticosteroids may depend not only on hormone availability or receptor expression, but also on the functional hierarchy between GR and MR, whereby signalling through one receptor may constrain or redirect the biological output of the other.

In mammals, circulating glucocorticoid concentrations rise during late gestation, coordinating the maturation of multiple organs, including the heart, in preparation for extrauterine life (Fowden et al. 1998; Bird et al. 2015). In the foetal myocardium, glucocorticoid-induced GR signalling contributes to cardiomyocyte structural, functional and metabolic maturation (Rog-Zielinska et al. 2013; Rog-Zielinska et al. 2015). Yet, prenatal studies have also reported divergent effects of cortisol on cardiomyocyte growth. In foetal sheep, sub-pressor intracoronary cortisol infusion increased cardiomyocyte cell-cycle activity without inducing hypertrophy or binucleation (Giraud et al. 2006), whereas high-dose amniotic cortisol infusion caused left ventricular cardiomyocyte hypertrophy without increasing cardiomyocyte number (Lumbers et al. 2005). Because these studies differed in dose, route, duration and haemodynamic context, the determinants of these divergent responses remain unclear. Subsequent pharmacological evidence suggested that cortisol-induced proliferation in the foetal heart is mediated, at least in part, through MR (Feng et al. 2013). These observations raise the possibility that MR may support a pro-proliferative corticosteroid response in cardiomyocytes, but whether this programme persists after birth, and how it is constrained by GR, remains unknown.

After birth, cardiomyocyte maturation coincides with a rapid loss of regenerative competence. Although transient cardiomyocyte proliferation supports heart regeneration during the neonatal period, this capacity is rapidly extinguished as cardiomyocytes withdraw from the cell cycle and acquire the structural, metabolic and functional features of mature adult cells, leaving the adult myocardium largely unable to replace cardiomyocytes lost after injury (Porrello et al. 2011; Uygur and Lee 2016; Bongiovanni et al. 2021). Defining the endogenous signals that impose this transition from regenerative growth to stable cell-cycle arrest therefore remains a central challenge in cardiovascular biology. We previously showed that corticosterone, the principal endogenous glucocorticoid in rodents, promotes cardiomyocyte cell-cycle exit and postnatal maturation through GR. Consistent with this, cardiomyocyte-specific GR ablation or transient pharmacological GR inhibition prolongs cardiomyocyte proliferative competence and enhances regenerative responses after myocardial infarction (Pianca et al. 2022). GR activation also limits the response of cardiomyocytes to regenerative growth factors and cytokines by inducing negative regulators of MAPK-ERK signalling (Da Pra et al. 2026). Together, these findings identify GR as a major endogenous brake on cardiomyocyte proliferation and cardiac regeneration.

By contrast, MR signalling in the adult heart has been studied predominantly in relation to aldosterone-dependent inflammation, fibrosis, hypertrophy, arrhythmogenesis and adverse remodelling (Gravez et al. 2013; Richardson et al. 2016; Ayuzawa and Fujita 2021). Cardiomyocyte-specific MR deletion does not produce an overt cardiac phenotype under basal conditions (Oakley et al. 2019) but protects against selected pathological responses after cardiac injury (Fraccarollo et al. 2011). Whether MR can regulate cardiomyocyte proliferative competence or contribute to cardiac regeneration is therefore unknown.

Beyond receptor hierarchy, an additional unresolved question is whether corticosteroid output is determined solely by receptor identity or also by the cellular context in which receptor activation occurs. Cardiomyocytes are the principal cellular effectors of myocardial regeneration, whereas fibroblasts and other stromal populations shape post-injury tissue remodelling and can directly modulate cardiomyocyte proliferative and regenerative competence through paracrine and extracellular-matrix-derived signals (Ieda et al. 2009; Wang et al. 2020). Because stromal MR activation is closely linked to fibrotic and inflammatory responses, activation of the same receptor in different cardiac compartments may generate divergent or even opposing tissue-level outcomes (Rickard and Young 2009; Rickard et al. 2012; Lother et al. 2019; Buffolo et al. 2022). Moreover, whether nuclear engagement of MR necessarily produces equivalent biological outputs under different ligand and receptor conditions remains unclear. Thus, how GR-MR hierarchy and cellular context are integrated to determine the cardiac response to endogenous corticosteroids remains unresolved.

Here, we show that endogenous corticosteroids are decoded in the heart through hierarchical and context-dependent GR-MR signalling. Genetic ablation or pharmacological inhibition of GR enables corticosterone to promote cardiomyocyte proliferation through MR, converting the output of the same physiological ligand from anti-proliferative to proliferative. Direct activation of MR by aldosterone further reveals that GR loss or inhibition enhances MR-driven proliferation in cardiomyocyte-enriched cultures, indicating that GR restrains the proliferative output of MR beyond simply controlling ligand allocation between the two receptors. However, this enhanced cardiomyocyte response is lost in mixed cardiac cultures, demonstrating that the surrounding cellular compartment gates the tissue-level consequences of MR activation.

We further show that the consequences of MR activation depend on ligand, receptor and cellular context: direct MR activation by aldosterone, in fact, stimulates stromal proliferation and fibroblast activation, whereas glucocorticoid-driven MR signalling under GR inhibition promotes cardiomyocyte proliferation without inducing a profibrotic stromal response. Notably, both aldosterone and corticosterone combined with GR inhibition promote MR nuclear localisation in cardiac fibroblasts, showing that their divergent stromal effects occur despite nuclear MR engagement under both conditions. Finally, corticosterone in the presence of GR antagonism promotes MR-dependent cardiomyocyte cell-cycle activity in adult murine myocardium, with analogous responses in porcine and human myocardium.

Together, our findings identify a functional GR-MR hierarchy, shaped by cellular context, as a mechanism of endocrine decoding in the heart and reveal how endogenous corticosteroid signalling can be redirected towards cardiomyocyte regenerative competence.

## RESULTS

### GR ablation uncovers a proliferative response to corticosterone

We previously showed that physiological glucocorticoids promote cardiomyocyte cell-cycle exit through activation of the glucocorticoid receptor (GR), thereby limiting the regenerative capacity of the neonatal heart (Pianca et al. 2022). We therefore asked whether endogenous corticosteroids retain any effect on cardiomyocyte proliferative activity once GR signalling is eliminated.

To address this question, we isolated primary neonatal cardiomyocytes from cardiomyocyte-specific GR conditional knockout (GR-cKO) mice and littermate controls (Pianca et al. 2022). Cardiomyocyte proliferation was assessed using complementary markers of cell-cycle progression, DNA synthesis and cytokinesis. As expected, corticosterone suppressed cardiomyocyte proliferation in control cultures. Unexpectedly, however, the same treatment increased the proportion of cycling cardiomyocytes in GR-deficient cultures (**Fig. 1a**). In cardiomyocytes, this response was also accompanied by increased BrdU incorporation (**Fig. 1b**) and a higher frequency of Aurora B-positive cleavage furrows (**Fig. 1c**), indicating enhanced DNA synthesis and progression through cytokinesis.

**Figure 1.**
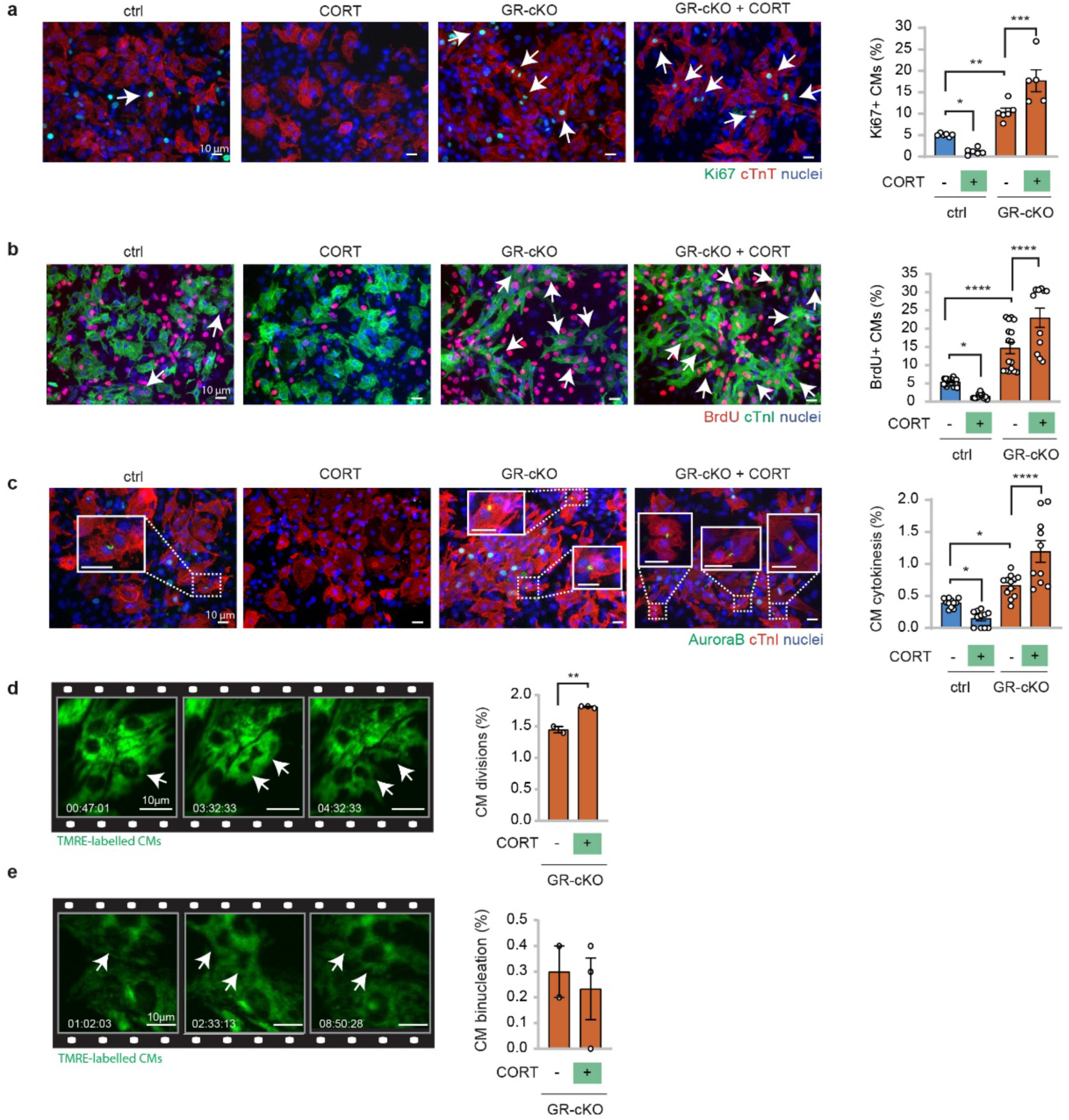
Physiological glucocorticoids promote proliferation of GR-deficient neonatal cardiomyocytes. (**a–c**) Cardiomyocytes (CMs) were isolated from postnatal day 1 (P1) GR^flox/flox^ control and cardiomyocyte-specific GR knockout (GR-cKO) mice, cultured *in vitro* and treated with corticosterone (CORT, 10 nM). CMs were identified by cTnI or cTnT immunostaining, as indicated in the Methods, and analysed for cell-cycle activity by Ki67 staining (**a**; n = 20 samples; 6686 CMs analysed), DNA synthesis by BrdU incorporation (**b**; n = 65 samples; 20569 CMs analysed) and cytokinesis by Aurora B kinase localization at the cleavage furrow (**c**; n = 45 samples; 22401 CMs analysed). Representative images are shown; arrows indicate proliferating CMs. Scale bars, 10 μm; (**d,e**) Quantification and representative frames from 24-h time-lapse imaging of TMRE-labelled CMs isolated from P1 GR-cKO mice and treated with vehicle or CORT (10 nM). Cardiomyocyte division events are shown in **d** and binucleation events in **e** (n = 5 samples; 2212 CMs analysed). Arrows indicate CMs undergoing the indicated cell-cycle events. Time is shown as hours:minutes. Scale bars, 10 μm; Data are presented as mean ± s.e.m. Statistical significance was assessed by one-way ANOVA followed by Šidák’s multiple-comparisons test in **a–c**, and by two-sided Student’s t-test in **d**, **e**. *P < 0.05, **P < 0.01, ***P < 0.001, ****P < 0.0001.

To determine whether these effects translate into productive cell division, we performed 24-hour time-lapse imaging of TMRE-labelled GR-deficient neonatal cardiomyocytes. Live-cell imaging directly confirmed that corticosterone increased cardiomyocyte division events (**Fig. 1d; Supplementary Video 1**) without significantly affecting the frequency of cardiomyocyte binucleation (**Fig. 1e**; **Supplementary Video 2**).

These findings reveal an unexpected property of endogenous corticosterone. Rather than being intrinsically anti-proliferative, corticosterone acquires proliferative activity when GR signalling is removed, suggesting that endogenous corticosteroids engage an alternative proliferative pathway normally masked by GR.

### Corticosterone extends the postnatal proliferative window in GR-deficient cardiomyocytes

We next asked whether the proliferative response to corticosterone observed in GR-deficient cardiomyocytes also occurs *in vivo* during postnatal heart development. Cardiomyocytes retain proliferative competence during the first days after birth, but this capacity is rapidly lost as the myocardium undergoes postnatal maturation. We therefore administered corticosterone to lactating mothers from postnatal day 0-1 to postnatal day 7 and analysed cardiomyocyte growth and cell-cycle activity in GR-cKO and control (GR^flox/flox^) pups (**Fig. 2a**).

**Figure 2.**
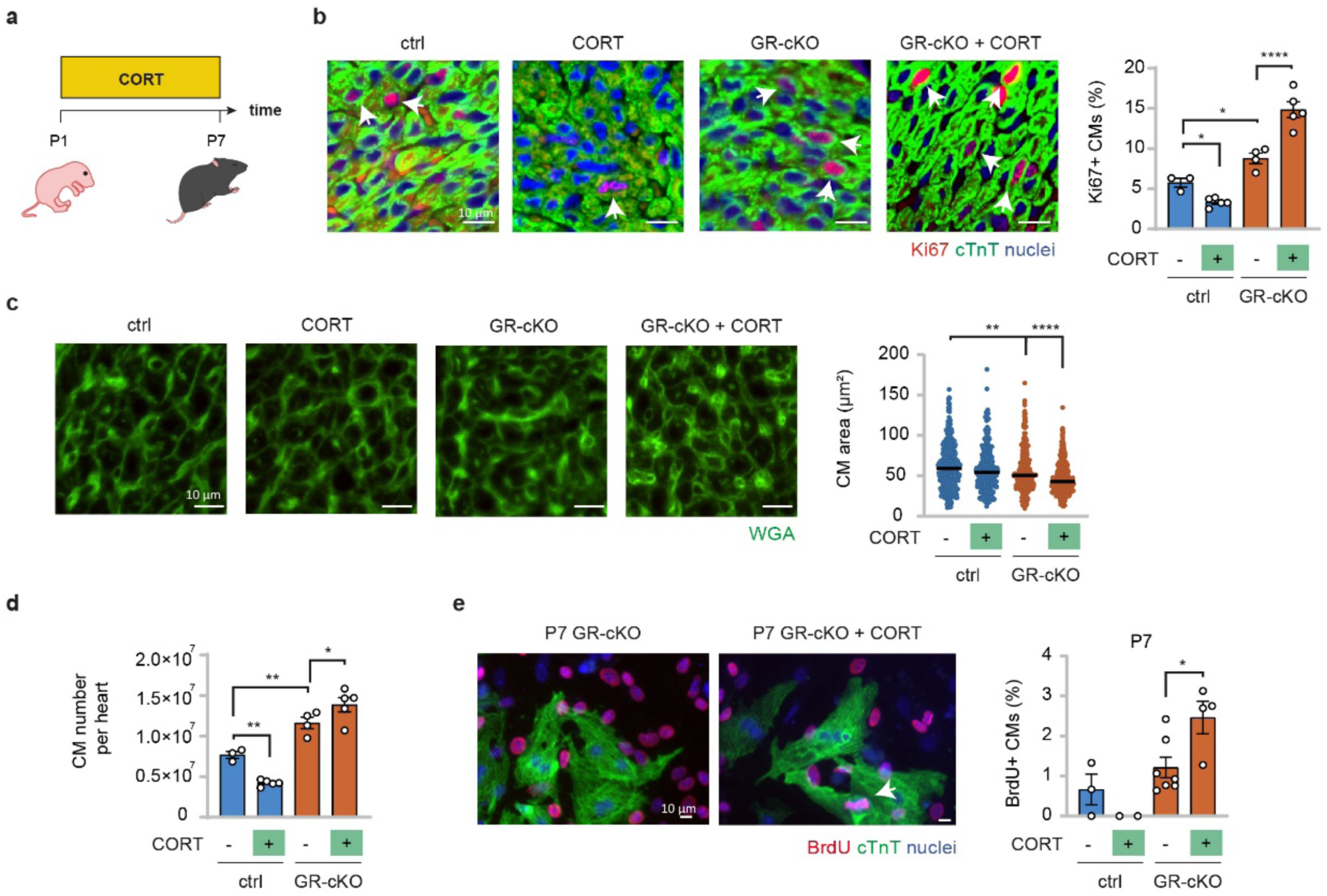
GR loss enables corticosterone to prolong the postnatal cardiomyocyte proliferative window. (**a–d**) Water-soluble corticosterone–HBC complex (100 µg/ml) was administered in the drinking water of lactating mothers to treat GR^flox/flox^ control and GR-cKO pups from postnatal day 0–1 (P0–P1) to postnatal day 7 (P7), as schematized in **a**. Cardiomyocyte cell-cycle activity was assessed in heart sections by Ki67 and cardiac troponin T immunostaining (**b**; n = 17 mice; 18693 CMs analysed). Cardiomyocyte cross-sectional area was measured by wheat germ agglutinin (WGA) staining (**c**; n = 17 mice; 1216 CMs analysed). Cardiomyocyte number was estimated by stereological analysis of P7 control and GR-cKO hearts after corticosterone treatment (**d**; n = 17 mice); (**e**) Immunofluorescence analysis of cardiomyocyte proliferation by BrdU incorporation in P7 control and GR-cKO cardiomyocyte cultures treated with vehicle or corticosterone (CORT, 10 nM) (n = 15 samples; 3095 CMs analysed). Representative images are shown; arrows indicate cycling CMs. Scale bars, 10 μm. Data are presented as mean ± s.e.m. in **b, d** and **e**, and as median in **c**. In all panels, statistical significance was assessed by one-way ANOVA followed by Šidák’s multiple-comparisons test. *P < 0.05, **P < 0.01, ***P < 0.001, ****P < 0.0001.

Corticosterone reduced body weight in both control and GR-cKO pups (**Supplementary Fig. 1a**), consistent with the known growth-restrictive effects of sustained glucocorticoid exposure during early postnatal life (Miyaso et al. 2022). In control pups, corticosterone reduced cardiomyocyte proliferation (**Fig. 2b**), without significantly affecting cardiomyocyte cross-sectional area (**Fig. 2c**), in agreement with the anti-proliferative effect of GR activation we previously assessed during the first days of postnatal life (Pianca et al. 2022; Da Pra et al. 2026). This reduction in cardiomyocyte proliferation was associated with decreased heart weight (**Supplementary Fig. 1b**).

In GR-cKO pups, corticosterone produced the opposite cardiac response. Despite showing reduced body weight, corticosterone-treated GR-cKO pups maintained heart weight comparable to untreated controls, resulting in an increased heart-weight-to-body-weight ratio (**Supplementary Figs. 1b, c**). This was accompanied by increased cardiomyocyte cell-cycle activity (**Fig. 2b**), reduced cardiomyocyte cross-sectional area (**Fig. 2c**) and a higher number of cardiomyocytes per heart estimated by stereological analysis (**Fig. 2d**). Thus, in the absence of cardiomyocyte GR, corticosterone shifts postnatal cardiomyocyte development away from hypertrophic maturation and towards hyperplastic growth.

To determine whether this response could be elicited beyond the early neonatal proliferative phase, we isolated cardiomyocytes from postnatal day 7 (P7) hearts, a stage at which cardiomyocytes have largely entered a post-mitotic state. Corticosterone increased proliferation in P7 GR-cKO cardiomyocytes (**Fig. 2e**), indicating that GR loss enables corticosterone to extend the physiological postnatal proliferative window. These findings show that corticosterone exerts opposite effects on cardiomyocyte proliferation depending on GR availability, both *in vitro* and during postnatal heart development *in vivo*. When GR is present, corticosterone favours cardiomyocyte cell-cycle exit and postnatal maturation; when cardiomyocyte GR is removed, the same hormone sustains proliferative competence during early heart development.

### MR mediates the proliferative output of corticosterone in the absence of GR

Because GR deletion was restricted to cardiomyocytes, we first examined whether corticosterone might indirectly promote their proliferation through GR activation in the surrounding stromal compartment. Because corticosterone can activate both GR and the mineralocorticoid receptor (MR), we employed dexamethasone, a synthetic glucocorticoid with strong GR-selective activity. Dexamethasone reduced cardiomyocyte proliferation in control cultures, consistent with the anti-proliferative activity of GR in cardiomyocytes (**Fig. 3a**). However, it did not reproduce the proliferative effect of corticosterone in GR-cKO cultures (**Fig. 3a**), arguing against an indirect mechanism mediated by GR activation in non-cardiomyocyte cells.

**Figure 3.**
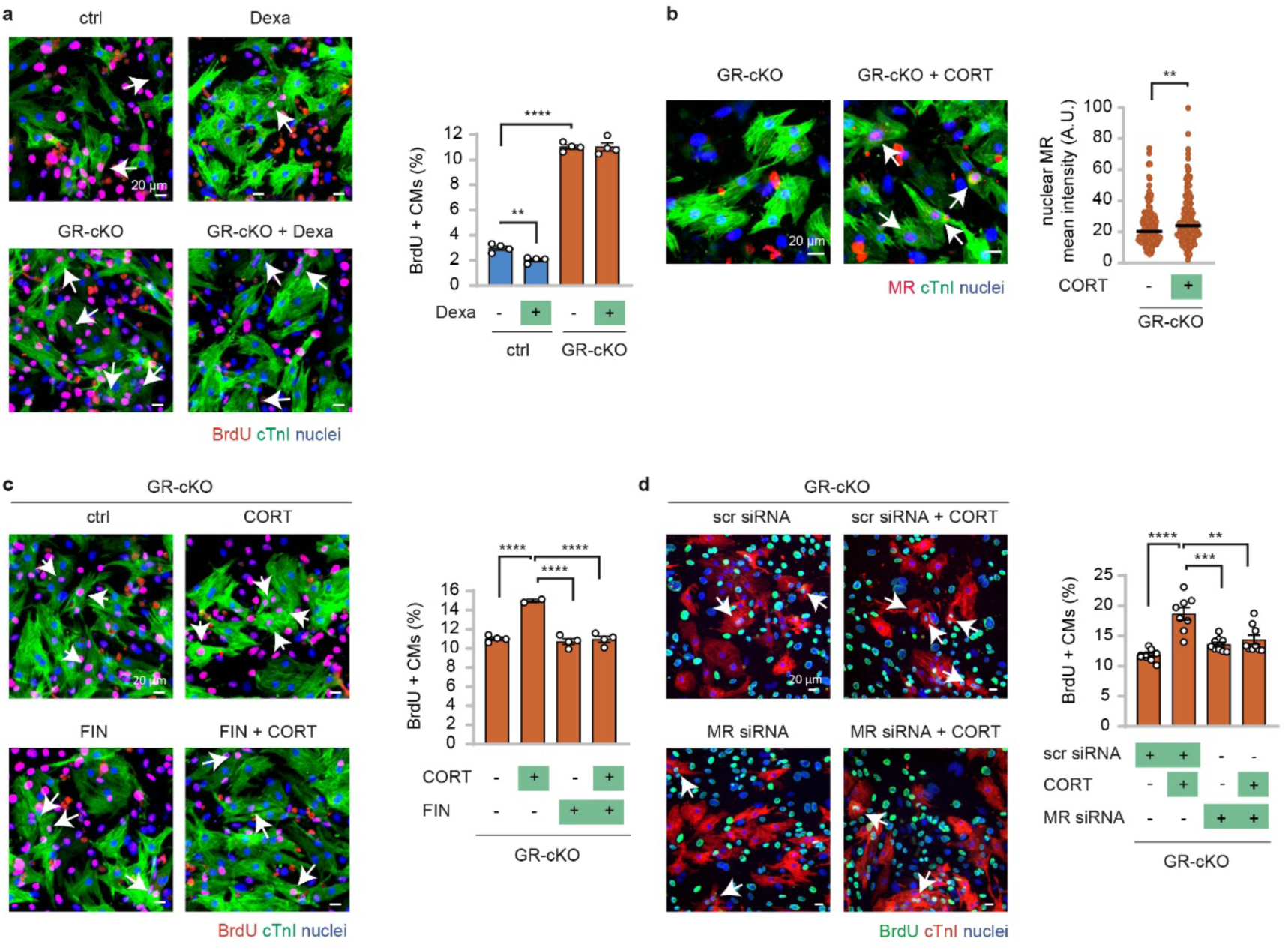
MR mediates the pro-proliferative response to corticosterone in GR-deficient cardiomyocytes. (**a**) Cardiomyocyte proliferation was assessed by BrdU incorporation in neonatal P1 cardiomyocytes isolated from GR^flox/flox^ control and GR-cKO hearts and treated with the GR-selective agonist dexamethasone (Dexa, 10 nM) (n = 16 samples; 3063 CMs analysed); (**b**) Nuclear MR mean fluorescence intensity was quantified by immunofluorescence in GR-cKO cardiomyocytes after short-term corticosterone exposure (CORT, 1 μM, 1 h) (375 CMs analysed in 6 samples); (**c**, **d**) P1 GR-cKO cardiomyocytes were treated with vehicle or corticosterone (CORT, 10 nM) in combination with either the MR antagonist finerenone (FIN, 100 nM) (**c**; n = 14 samples; 2989 CMs analysed) or MR-specific siRNA (**d**; n = 36 samples; 14649 CMs analysed). Scrambled siRNA (scr) was used as control. Representative images are shown; arrows indicate proliferating CMs. Scale bars, 20 μm; Data are presented as mean ± s.e.m. in **a**, **c** and **d**, and as median in **b**. Statistical significance was assessed by one-way ANOVA followed by Šidák’s multiple-comparisons test in **a**, **c** and **d**, and by two-sided Student’s t-test in **b**. *P < 0.05, **P < 0.01, ***P < 0.001, ****P < 0.0001.

We next considered MR as the alternative corticosteroid receptor responsible for the corticosterone-induced proliferative response. MR binds physiological glucocorticoids with high affinity and, because the glucocorticoid-inactivating enzyme 11β-hydroxysteroid dehydrogenase type 2 is minimally expressed in the heart, cardiac MR can be activated by circulating glucocorticoids (Richardson et al. 2016). Since ligand binding promotes MR nuclear translocation, we assessed its nuclear localisation following corticosterone treatment as a readout of receptor activation. Corticosterone effectively increased nuclear MR signal in GR-deficient cardiomyocytes (**Fig. 3b**). To establish whether MR was required for the corticosterone-driven proliferative response, we treated GR-cKO cultures with finerenone, a selective non-steroidal MR antagonist. Finerenone abolished the corticosterone-induced increase in cardiomyocyte proliferation (**Fig. 3c**). We independently confirmed this finding by silencing MR using a specific MR-targeting siRNA. Efficient reduction of MR expression was verified by quantitative gene-expression analysis (**Supplementary Fig. 2a**), and MR knockdown markedly attenuated corticosterone-induced cardiomyocyte proliferation in GR-cKO cultures (**Fig. 3d**).

To assess the specificity of this response, we individually silenced the other steroid hormone receptors, namely androgen receptor (AR), oestrogen receptor-α (ERα), oestrogen receptor-β (ERβ), G protein-coupled oestrogen receptor 1 (GPER1) and progesterone receptor (PR), confirming the efficient reduction in their gene expression level (**Supplementary Figs. 2b-f**). None of these interventions prevented the proliferative effect of corticosterone (**Supplementary Fig. 2g**), identifying MR as the steroid receptor specifically required for this response.

Together, these findings demonstrate that GR loss does not merely remove an anti-proliferative pathway. Instead, it changes how corticosterone is decoded, allowing the same physiological ligand to promote cardiomyocyte proliferation through MR. The biological output of glucocorticoids therefore depends on the functional hierarchy between GR and MR, with GR exerting a dominant effect under physiological conditions, rather than on ligand identity alone.

### Stromal context determines the biological output of MR signalling

The finding that MR mediates the proliferative effect of corticosterone in GR-deficient cardiomyocytes raised an apparent paradox. In the adult heart, MR activation has been primarily associated with fibrosis, inflammation and adverse remodelling rather than with regeneration. We therefore asked whether direct MR activation by aldosterone could reproduce the proliferative response elicited by corticosterone in the absence of GR.

To activate MR without relying on the redistribution of corticosterone between GR and MR, we treated mixed neonatal cardiac cultures with aldosterone, the principal endogenous mineralocorticoid. In these multicellular cultures, aldosterone failed to increase cardiomyocyte proliferation (**Fig. 4a**), despite the ability of MR to promote cardiomyocyte proliferation when engaged by corticosterone in the absence of GR. However, aldosterone promoted the proliferation of stromal cells cultured in this multicellular context (**Fig. 4a**). Notably, aldosterone remained unable to increase cardiomyocyte proliferation in mixed cultures when GR was genetically ablated in cardiomyocytes or pharmacologically antagonised with RU486 (**Figs. 4b, c**). These findings raised the possibility that the biological output of MR activation is shaped not only by ligand identity, but also by the cellular context in which the receptor is engaged. To test this possibility, we separated cardiomyocyte-enriched and stromal-enriched cardiac cultures. In cardiomyocyte-enriched cultures, aldosterone increased cardiomyocyte proliferation (**Fig. 4d**), indicating that direct MR activation is sufficient to promote cardiomyocyte cell-cycle activity when stromal influence is minimized. This response was further enhanced in GR-cKO cardiomyocytes compared to control ones, and was similarly increased by pharmacological GR antagonism (**Figs. 4e, f**). These findings indicate that GR restrains the intrinsic proliferative output of MR in cardiomyocytes even when MR is directly stimulated, and therefore that the effect of GR loss cannot be explained exclusively by redistribution of corticosterone between GR and MR. In stromal-enriched cultures, aldosterone also stimulated fibroblast proliferation (**Fig. 4g**) and increased the proportion of activated fibroblasts (**Fig. 4h**), as assessed by α-smooth muscle actin staining. Thus, aldosterone engages MR and stimulates biological programmes in both cardiomyocytes and stromal cells, inducing cardiomyocyte proliferation when stromal influence is minimised, while promoting fibroblast expansion and activation in the stromal compartment.

**Figure 4.**
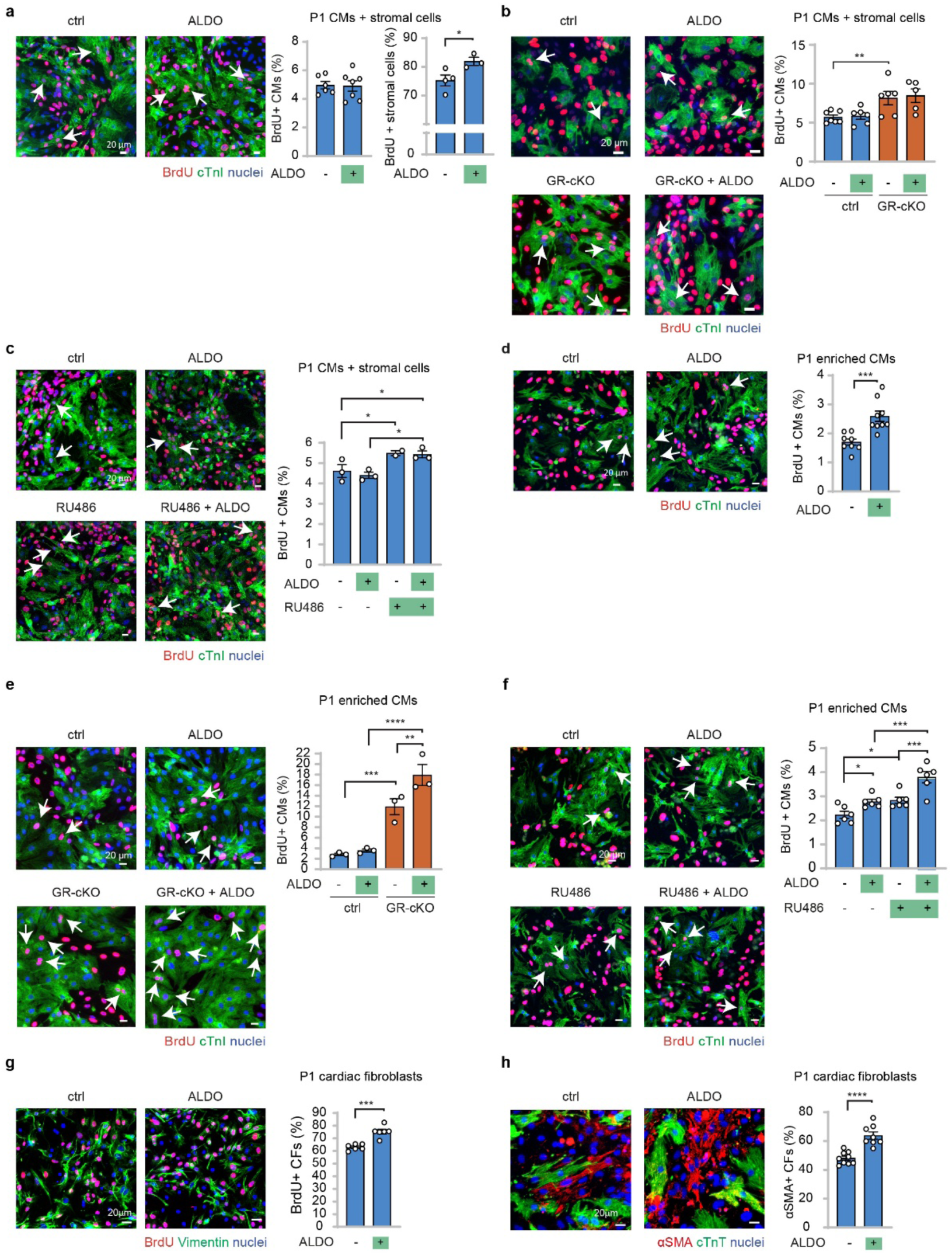
Stromal context determines the biological output of aldosterone-driven MR activation. (**a**) Primary neonatal cardiac cells isolated from postnatal day 1 (P1) mice were treated with vehicle or MR agonist aldosterone (ALDO, 10 nM), and BrdU incorporation was quantified in cardiomyocytes (left) and cardiac stromal cells (right) (n = 13 samples with 4018 CMs analysed; n = 8 samples with 2696 stromal cells analysed); (**b**) P1 cardiac cells were isolated from control (ctrl) and cardiomyocyte-specific GR ablated (GR-cKO) hearts and treated with ALDO (10 nM) (n = 24 samples with 7525 CMs analysed); (**c**) Cardiac cells isolated from P1 mice were administered with ALDO (10 nM) and/or RU486 (100 nM) (n = 11 samples with 2003 CMs analysed); (**d**) P1 cardiac cell cultures were enriched for cardiomyocytes and treated with vehicle or ALDO (10 nM). BrdU incorporation was quantified in troponin-positive cardiomyocytes (n = 17 samples; 5580 CMs analysed); (**e**) P1 cardiac cells were isolated from ctrl and GR-cKO hearts, enriched for cardiomyocytes and treated with ALDO (10 nM) (n = 12 samples with 3449 CMs analysed); (**f**) Cardiac cells were isolated from P1 mice, enriched for cardiomyocytes and administered with ALDO (10 nM) and/or RU486 (100 nM) (n = 24 samples with 6001 CMs analysed); (**g**) P1 cardiac cell cultures were enriched for cardiac stromal cells and treated with vehicle or ALDO (10 nM). BrdU incorporation was quantified in vimentin-positive stromal cells (n = 12 samples; 4482 stromal cells analysed); (**h**) Fibroblast activation was assessed after ALDO treatment (10 nM) by α-smooth muscle actin (αSMA) and cardiac troponin T (TnT) immunostaining. αSMA positivity was quantified in TnT-negative cells (n = 17 samples; 6711 cells analysed). Representative images are shown. Scale bars, 20 μm. Data are presented as mean ± s.e.m. Statistical significance was assessed by one-way ANOVA followed by Šidák’s multiple-comparisons test in **b**, **c**, **e** and **f**, and by two-sided Student’s t-test in **a**, **d**, **g** and **h**. *P < 0.05, **P < 0.01, ***P < 0.001, ****P < 0.0001.

Together, these findings identify two levels of control over the proliferative response to MR activation. GR imposes a cardiomyocyte-intrinsic restraint on MR-driven proliferation, whereas the stromal compartment prevents this enhanced intrinsic response from emerging in multicellular cardiac contexts. MR signalling is therefore not intrinsically regenerative or profibrotic. Rather, its tissue-level output is determined by the functional hierarchy between GR and MR within cardiomyocytes and by the surrounding cellular context.

### GR inhibition redirects endogenous corticosteroid signalling towards a cardiomyocyte-selective proliferative response

Direct MR activation by aldosterone promoted cardiomyocyte proliferation in cardiomyocyte-enriched cultures, and this response was further enhanced by pharmacological or genetic disruption of GR. However, neither aldosterone alone nor aldosterone combined with GR inhibition increased cardiomyocyte proliferation in mixed cardiac cultures, indicating that removal of the GR-dependent brake is not sufficient to generate a net proliferative response in the presence of the stromal compartment. We therefore asked whether redirecting physiological glucocorticoids towards MR could produce a distinct, more cardiomyocyte-selective output in a multicellular cardiac context.

In mixed neonatal cardiac cultures, corticosterone alone reduced cardiomyocyte proliferation (**Fig. 5a**), consistent with the dominant anti-proliferative effect of GR signalling (Pianca et al. 2022). By contrast, combined treatment with corticosterone and the GR antagonist RU486 increased cardiomyocyte proliferation in the same multicellular context (**Fig. 5a**). Importantly, this response occurred without concomitantly increasing stromal cell proliferation above control levels (**Fig. 5a**), suggesting that GR inhibition redirects corticosterone towards a proliferative cardiomyocyte output without broadly stimulating non-myocyte expansion. The ability of corticosterone combined with GR antagonist RU486, but not aldosterone combined with RU486, to promote cardiomyocyte proliferation in mixed cultures further indicates that GR antagonism alone does not account for the different tissue-level responses.

**Figure 5.**
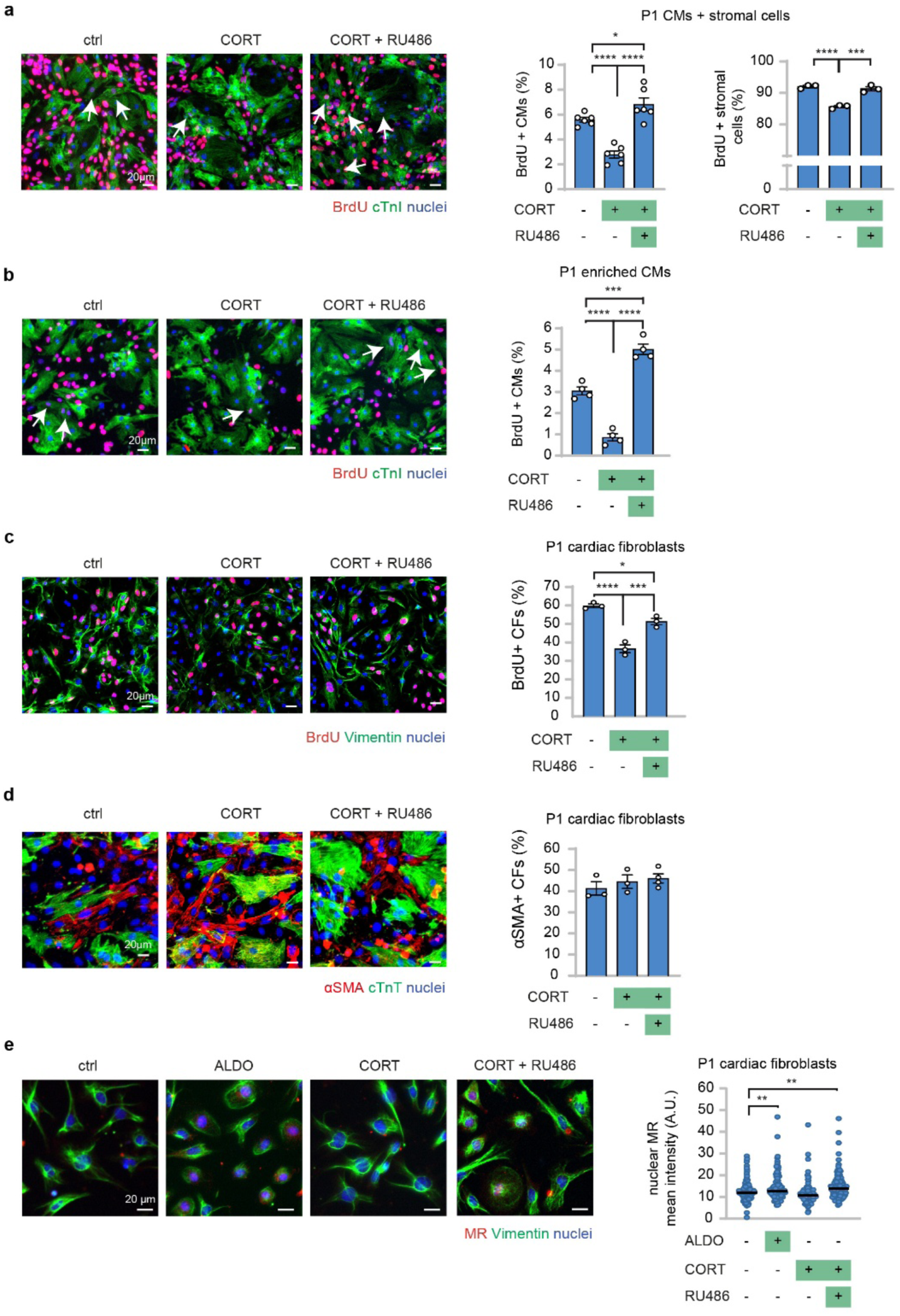
GR inhibition redirects corticosterone signalling towards a cardiomyocyte-selective proliferative response. (**a**) Primary neonatal cardiac cell cultures were treated with vehicle, corticosterone (CORT, 10 nM), or CORT plus RU486 (100 nM), and BrdU incorporation was quantified in cardiomyocytes (left) and cardiac stromal cells (right) (n = 18 samples with 4124 CMs analysed; n = 9 samples with 2465 stromal cells analysed); (**b, c**) Cardiomyocyte-enriched (**b**) and stromal-enriched (**c**) neonatal cardiac cultures were treated with vehicle, CORT (10 nM), or CORT plus RU486 (100 nM). BrdU incorporation was quantified in troponin-positive cardiomyocytes (**b**; n = 12 samples; 3475 CMs analysed) or vimentin-positive stromal cells (**c**; n = 9 samples; 2696 stromal cells analysed); (**d**) Fibroblast activation was assessed in neonatal primary cardiac cell cultures treated with vehicle, CORT (10 nM), or CORT plus RU486 (100 nM). αSMA positivity was quantified in TnT-negative cells (n = 10 samples; 4511 cells analysed); (**e**) Nuclear MR mean fluorescence intensity was quantified by immunofluorescence in cardiac fibroblasts after 1 hour exposure to aldosterone (ALDO, 1 μM), corticosterone (CORT, 1 μM) or CORT plus RU486 (10 μM) (467 cells analysed in 12 samples). Representative images are shown. Scale bars, 20 μm. Data are presented as mean ± s.e.m. in panels **a-d** and as median in **e**. In all panels, statistical significance was assessed by one-way ANOVA followed by Šidák’s multiple-comparisons test. *P < 0.05, **P < 0.01, ***P < 0.001, ****P < 0.0001.

We then examined cardiomyocyte-enriched and stromal-enriched cultures separately. In cardiomyocyte-enriched cultures, corticosterone combined with GR antagonist RU486 increased cardiomyocyte proliferation (**Fig. 5b**), showing that the combined removal of GR signalling and exposure to corticosterone is sufficient to promote cardiomyocyte cell-cycle activity when stromal influence is minimised. In stromal-enriched cultures, however, the same treatment did not increase fibroblast proliferation (**Fig. 5c**). Moreover, corticosterone, either alone or in combination with GR antagonist RU486, failed to induce fibroblast activation, as assessed by α-smooth muscle actin staining (**Fig. 5d**). To determine whether the lack of a fibroblast response reflected insufficient MR engagement, we assessed MR subcellular localisation in cardiac fibroblasts. Both aldosterone and combined corticosterone-RU486 treatment increased MR nuclear localisation compared with vehicle, indicating that both conditions promote MR nuclear engagement in the stromal compartment (**Fig. 5e**).

Thus, the biological consequences of MR engagement differ according to how the receptor is activated. Direct stimulation by aldosterone activates both cardiomyocyte and stromal programmes, including fibroblast proliferation and activation. By contrast, corticosterone under GR inhibition promotes cardiomyocyte proliferation without triggering the profibrotic stromal response associated with aldosterone. Importantly, this divergence occurs despite nuclear localisation of MR in fibroblasts under both conditions, demonstrating that MR nuclear translocation is not sufficient to determine a uniform stromal output. Together with the finding that GR inhibition potentiates aldosterone-induced proliferation only when stromal influence is minimised, these results indicate that the cardiomyocyte selectivity of corticosterone plus GR antagonist RU486 cannot be explained by GR blockade or MR activation alone. Rather, it emerges from the combined effects of ligand context, hierarchical GR-MR signalling and cellular composition. Redirecting endogenous corticosteroid signalling may therefore provide a more selective route to enhance cardiomyocyte proliferative competence than direct mineralocorticoid stimulation.

### Myocardial injury enhances corticosteroid availability and cardiomyocyte MR nuclear engagement after GR inhibition

Having established that the functional hierarchy between GR and MR determines the proliferative output of corticosterone in neonatal cardiomyocytes, we next asked whether this mechanism could be engaged in the adult injured heart. Myocardial infarction is accompanied by a systemic stress response that increases circulating glucocorticoids, potentially providing endogenous ligand availability for corticosteroid receptor signalling.

We therefore measured serum corticosterone levels in adult mice after myocardial infarction. Circulating corticosterone was markedly increased after injury compared with non-infarcted animals in both control and GR-cKO mice (**Figs**. **6a****, b**), indicating that the infarcted heart is exposed to elevated endogenous corticosteroid levels. We then assessed whether this hormonal environment was associated with MR activation when GR signalling was genetically or pharmacologically inhibited.

**Figure 6.**
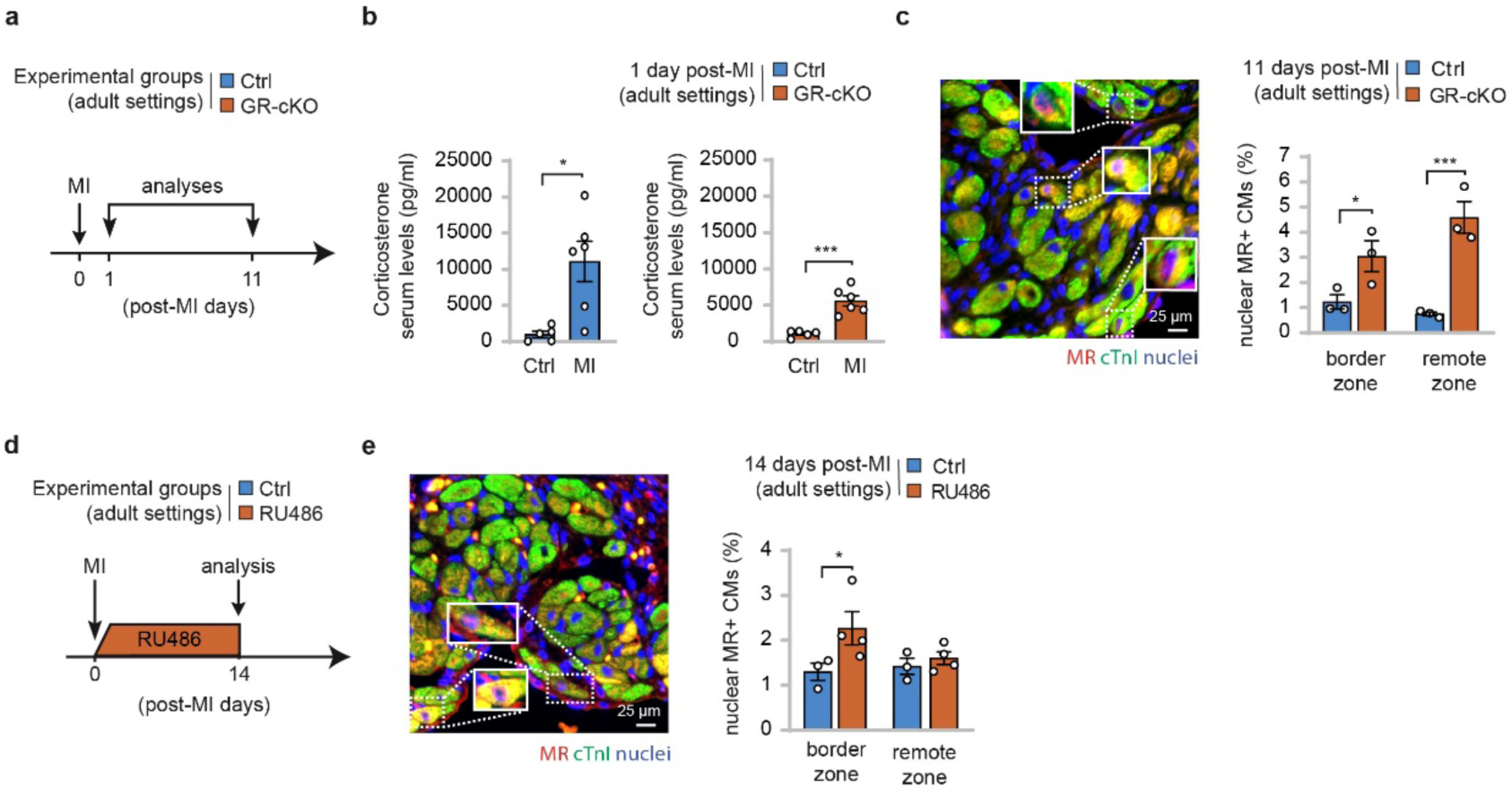
Increased corticosteroid availability after myocardial infarction enhances MR activation upon GR deletion or antagonisation. (**a**) Experimental scheme for analysis of control (Ctrl) and cardiomyocyte-specific GR knockout (GR-cKO) adult mice under basal conditions or after myocardial infarction, with analysis at the indicated time points; (**b**) Serum corticosterone levels were measured in Ctrl and GR-cKO adult mice under basal conditions or 24 h after myocardial infarction. Data are shown as two independent comparisons: basal versus infarcted Ctrl mice and basal versus infarcted GR-cKO mice (n = 11 mice per comparison); (**c**) MR nuclear localisation was assessed by MR and cardiac troponin I (cTnI) co-immunostaining in cardiomyocytes from the border and remote zones of infarcted Ctrl and GR-cKO hearts (n = 6 samples; 8261 CMs analysed); (**d**) Experimental scheme for analysis of adult mice subjected to myocardial infarction and treated with vehicle or the GR antagonist RU486 before analysis at the indicated time points after surgery; (**e**) MR nuclear localisation was assessed by MR and cTnI co-immunostaining in cardiomyocytes from the border and remote zones of infarcted vehicle- and RU486-treated hearts (n = 7 samples; 10948 CMs analysed). Representative images are shown. Scale bars, 25 μm. Data are presented as mean ± s.e.m. Statistical significance was assessed by two-sided Student’s t-test in **b**, **c** and **e**. *P < 0.05, **P < 0.01, ***P < 0.001, ****P < 0.0001.

MR activation was evaluated by quantifying its nuclear localisation in cardiomyocytes, identified by cardiac troponin I immunostaining. In adult cardiomyocyte-specific GR-cKO mice, myocardial infarction increased cardiomyocyte nuclear MR signal in both border and remote myocardium (**Fig. 6c**). Similarly, pharmacological GR antagonism after myocardial infarction increased cardiomyocyte MR nuclear localisation compared with vehicle-treated infarcted controls (**Figs. 6d, e**).

These findings indicate that myocardial injury generates an endocrine context in which endogenous corticosteroid levels are elevated, and that inhibition of GR enhances cardiomyocyte MR activation in the adult heart. Thus, the corticosteroid receptor redirection identified in neonatal cardiomyocytes can also be engaged under pathophysiological conditions relevant to cardiac injury. Importantly, we previously demonstrated that GR ablation or antagonization enhances cardiac regeneration and improves heart function following myocardial infarction (Pianca et al. 2022). The present findings raise the possibility that enhanced MR engagement contributes to these beneficial effects.

### Redirecting corticosteroid signalling promotes cardiomyocyte cell-cycle activity in adult mammalian myocardium

To determine whether receptor redirection of corticosteroid signalling can stimulate cardiomyocyte cell-cycle activity in adult myocardial tissue, we used living ventricular myocardial slices, which preserve the multicellular composition, extracellular matrix organization and tissue architecture of the adult heart (Watson et al. 2019; Caliandro et al. 2025; Husetić et al. 2026). This *ex vivo* system allowed us to test the effect of corticosterone and GR antagonism directly in mature myocardium while maintaining native cardiomyocyte-stromal interactions.

We first tested whether the GR-MR hierarchy identified in neonatal cardiomyocytes remains functionally operative in uninjured adult murine myocardium. Corticosterone reduced cardiomyocyte cell-cycle activity, whereas combined corticosterone and RU486 treatment increased the proportion of cycling cardiomyocytes (**Fig. 7a**). MR antagonist finerenone alone had no detectable effect, but abolished the increase induced by corticosterone plus RU486 (**Fig. 7a**), demonstrating that the response to corticosterone under GR antagonism requires MR also in mature myocardial tissue.

**Figure 7.**
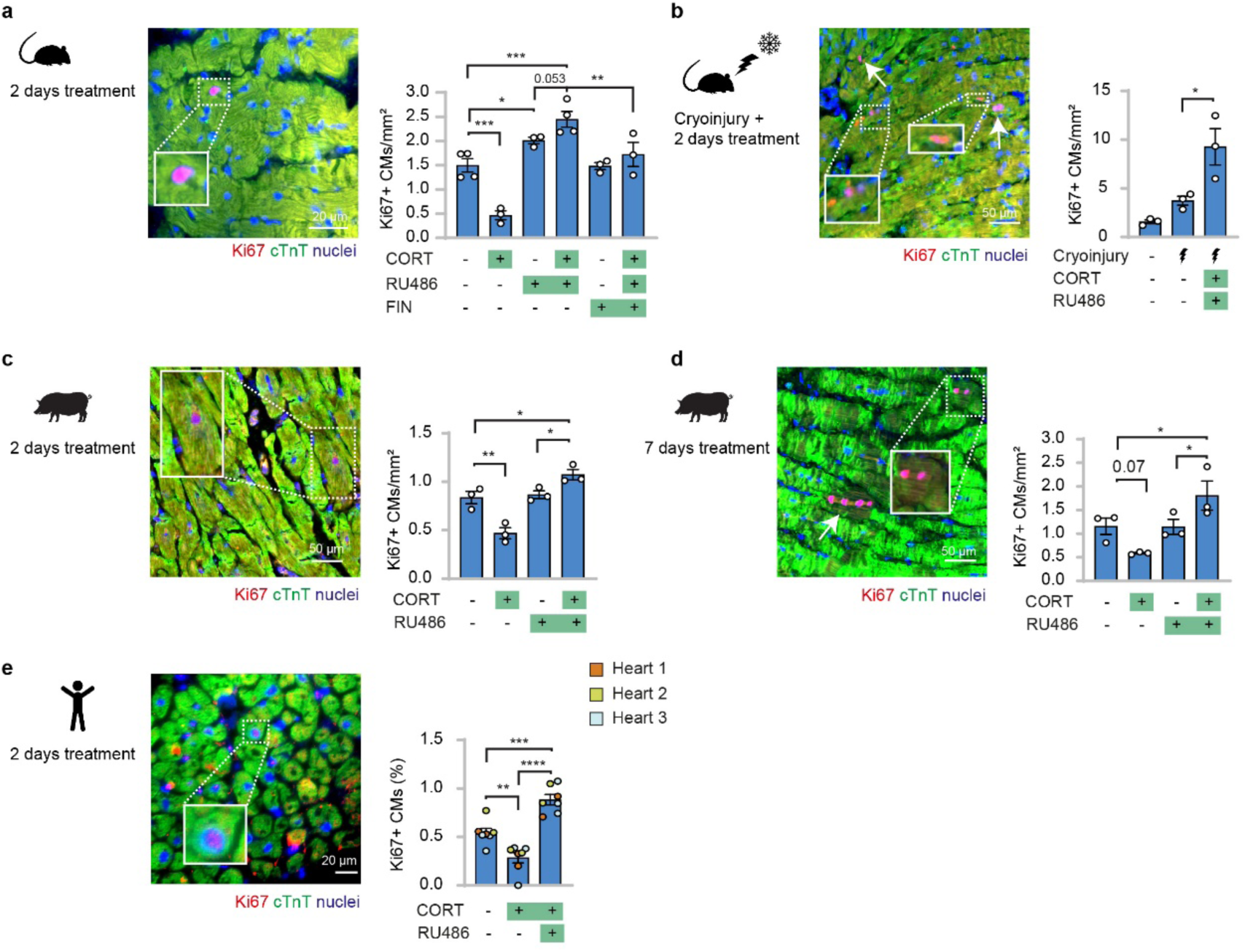
GR inhibition redirects corticosteroid signalling towards cardiomyocyte cell-cycle activity in adult myocardial tissue. (**a, b**) Living myocardial slices were generated from adult murine left ventricular tissue and (**a**) treated *ex vivo* with vehicle, corticosterone (CORT, 1 μM), RU486 (10 μM), finerenone (FIN, 10 μM) or the indicated combinations for 2 days (n = 20 samples) or (**b**) subjected to cryoinjury and cultured *ex vivo* for 2 days in medium containing vehicle, or CORT (1 μM) plus RU486 (10 μM) (n = 9 samples). Cardiomyocyte cell-cycle activity was assessed by Ki67 immunostaining. Representative images are shown. Scale bars, 20 μm (**a**) and 50 μm (**b**). (**c, d**) Adult swine myocardial slices were treated *ex vivo* with vehicle, corticosterone (CORT, 1 μM), RU486 (10 μM), or CORT plus RU486, and cardiomyocyte cell-cycle activity was assessed by Ki67 immunofluorescence after 2 days (**c**, n = 12 samples) or 7 days of culture (**d**, n = 12 samples). Representative images are shown. Scale bars, 50 μm. (**e**) Human left ventricular myocardial slices obtained from donor hearts were treated *ex vivo* with vehicle, CORT (1 μM), or CORT plus RU486 (10 μM) for 2 days. Cardiomyocyte cell-cycle activity was assessed by Ki67 immunofluorescence (n = 21 samples from 3 donors; 11513 CMs analysed). A representative image is shown. Scale bar, 20 μm. Data are presented as mean ± s.e.m. In all panels, statistical significance was assessed by one-way ANOVA followed by Šidák’s multiple-comparisons test. *P < 0.05, **P < 0.01, ***P < 0.001, ****P < 0.0001.

We next tested whether this response could also be engaged after myocardial injury. Murine left ventricular slices were subjected to cryoinjury, as previously described (Abbas et al. 2024), and treated *ex vivo* with corticosterone and GR antagonist RU486. Combined treatment further increased cardiomyocyte cell-cycle activity in the injured myocardium (**Fig. 7b**), supporting the ability of corticosteroid receptor redirection to potentiate the proliferative response after cardiac damage.

Having established MR dependence in adult murine myocardium, we next examined whether the response to corticosteroid receptor redirection is conserved in a large-mammal model. In adult porcine myocardial slices, corticosterone alone reduced cardiomyocyte cell-cycle activity, consistent with the anti-proliferative dominance of GR signalling observed in neonatal cultures. In contrast, combined treatment with corticosterone and GR antagonist RU486 increased the proportion of cycling cardiomyocytes after both 2 days and 7 days of culture (**Figs. 7c, d**). These findings indicate that receptor redirection of corticosteroid signalling can overcome the anti-proliferative effect of glucocorticoids in adult large-mammal myocardium.

Finally, we examined whether this response is conserved in human myocardium. Adult human left ventricular slices obtained from donor hearts were treated with corticosterone alone or in combination with GR antagonist RU486. As in porcine tissue, corticosterone alone reduced cardiomyocyte cell-cycle activity, whereas combined glucocorticoid treatment and GR antagonism increased the proportion of cycling cardiomyocytes (**Fig. 7e**).

Together, these data show that the functional GR-MR hierarchy identified in neonatal cardiomyocytes remains operative in adult murine myocardium, where the response to corticosterone under GR antagonism requires MR. The analogous responses to corticosterone plus GR antagonism observed in porcine and human myocardial slices support conservation of this corticosteroid response across adult mammalian species. Together, these findings support a model in which corticosteroid output is jointly determined by functional GR-MR hierarchy, ligand identity and cellular composition (**Fig. 8**).

**Figure 8.**
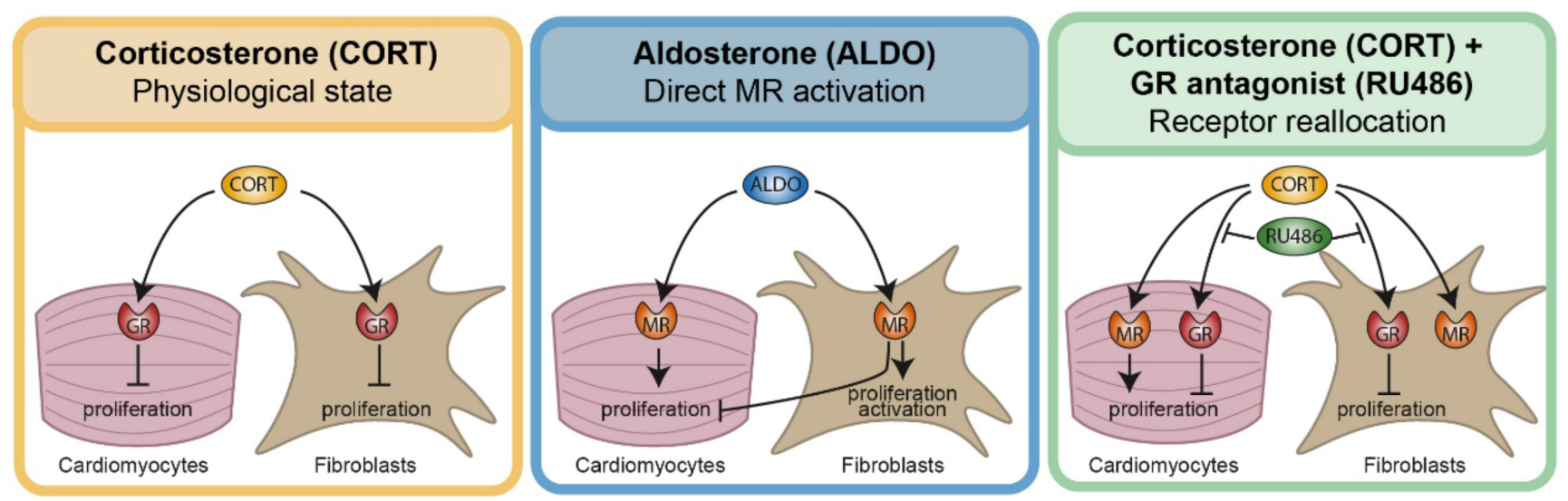
Model of receptor- and context-dependent corticosteroid decoding in the heart. Schematic model summarizing how corticosteroid receptor hierarchy and cellular context determine the biological output of corticosteroid signalling in cardiomyocytes and cardiac stromal cells. Under physiological receptor balance, corticosterone (CORT) signalling through the glucocorticoid receptor (GR) suppresses cardiomyocyte and cardiac fibroblast proliferation. On the other hand, direct mineralocorticoid receptor (MR) activation by aldosterone (ALDO) promotes cardiomyocyte mitogenic capacity. However, ALDO also stimulates cardiac stromal cell proliferation and fibroblast activation, and this stromal response is associated with loss of the cardiomyocyte pro-proliferative effect. Thus, ALDO-driven MR activation produces divergent cell-type-specific outputs, with stromal activation limiting the net proliferative response of cardiomyocytes in a multicellular setting. When GR is pharmacologically inhibited by RU486, CORT signalling is redirected towards MR, promoting cardiomyocyte proliferative competence without inducing stromal proliferation or profibrotic activation, resulting in a net increase of cardiomyocyte proliferation in the myocardial tissue. Together, these findings indicate that endogenous corticosteroids are decoded in the heart according to receptor hierarchy and stromal context, allowing corticosteroid signalling to favour either cardiomyocyte cell-cycle exit or proliferative competence.

## DISCUSSION

Endogenous corticosteroids are generally viewed as hormonal cues that promote cardiomyocyte maturation and restrict regenerative competence. Our findings show that this output is not an intrinsic property of the ligand, but depends on the receptor hierarchy through which it is decoded.

We previously showed that glucocorticoid-GR signalling promotes neonatal cardiomyocyte maturation, limits cardiomyocyte proliferative competence and constrains regenerative responses after myocardial infarction (Pianca et al. 2022), while attenuating cardiomyocyte responsiveness to regenerative growth factors and cytokines through induction of negative regulators of MAPK-ERK signalling (Da Pra et al. 2026). The present study identifies an additional property of GR: beyond directly imposing maturation and anti-proliferative programmes, GR is functionally dominant over MR and restrains the emergence of an alternative corticosteroid response. In fact, when GR is genetically removed or pharmacologically antagonised, corticosterone instead elicits an MR-dependent proliferative response. Disruption of GR also potentiates aldosterone-induced cardiomyocyte proliferation, demonstrating that GR restrains MR output beyond simply controlling ligand allocation between the two receptors. Endogenous corticosteroids are therefore not intrinsically anti-proliferative in the heart; their biological output is determined by functional GR-MR hierarchy.

A central implication of these findings is that MR signalling in the heart cannot be understood solely through the lens of aldosterone biology. In classical epithelial tissues, aldosterone is the dominant physiological MR ligand because 11β-hydroxysteroid dehydrogenase type 2 (11β-HSD2) protects MR from glucocorticoid occupancy. In the heart, however, low 11β-HSD2 expression allows endogenous glucocorticoids to access MR, placing the functional relationship between GR and MR at the centre of cardiac corticosteroid decoding (Richardson et al. 2016). This concept is consistent with the high affinity of MR for physiological glucocorticoids and with the established complexity of corticosteroid receptor selectivity (Hultman et al. 2005; Mifsud and Reul 2016). Our data show that corticosterone promotes cardiomyocyte proliferation through MR when GR is absent or inhibited, shifting the mechanistic question from which endogenous hormone can bind cardiac MR to how a shared corticosteroid input is interpreted according to the functional state of the two receptors.

The comparison with aldosterone further shows that GR dominance cannot be explained solely by competition for corticosterone. Direct activation of MR by aldosterone promoted cardiomyocyte proliferation in cardiomyocyte-enriched cultures, and this response was enhanced by either pharmacological GR antagonism or cardiomyocyte-specific GR ablation. GR therefore imposes a cardiomyocyte-intrinsic restraint on MR-dependent proliferation even when MR is activated independently of glucocorticoid redistribution. Previous evidence that cardiomyocyte-specific MR deletion modifies the cardiac consequences of concomitant GR ablation is consistent with functional interaction between these receptors (Oakley et al. 2019). Together, these observations support a hierarchical model in which GR shapes the biological output of MR at a level downstream of ligand availability.

Cellular context provides a second level of control. Aldosterone did not increase cardiomyocyte proliferation in mixed cardiac cultures, even when GR was genetically ablated in cardiomyocytes or pharmacologically antagonised, whereas a proliferative response emerged when stromal influence was reduced. In parallel, aldosterone stimulated fibroblast proliferation and activation. By contrast, corticosterone combined with GR inhibition promoted cardiomyocyte proliferation in mixed cultures without increasing stromal proliferation or myofibroblast activation. MR engagement, therefore, is not intrinsically regenerative or pathological; its tissue-level consequences depend on ligand identity, GR availability and the cellular compartment in which the response is generated.

The presence of the stromal compartment was associated with loss of the aldosterone-induced cardiomyocyte response observed in cardiomyocyte-enriched cultures, although the mechanism responsible for this context dependence remains unclear. Inhibitory paracrine signalling, altered ligand availability, extracellular-matrix-dependent cues or other multicellular interactions may contribute. Fibroblasts and other non-myocyte populations not only regulate extracellular matrix remodelling, inflammation and tissue mechanics after injury, but can also directly promote or restrain cardiomyocyte proliferative competence through paracrine and matrix-derived signals (Ieda et al. 2009; Wang et al. 2020). MR activation in non-myocyte compartments also contributes to pathological cardiac remodelling (Rickard and Young 2009; Lother et al. 2019; Buffolo et al. 2022). Defining whether stromal cells modify the cardiomyocyte response to aldosterone through secreted factors, extracellular-matrix cues or other multicellular mechanisms will be important for understanding how MR-dependent programmes are integrated at tissue level.

The contrasting fibroblast responses to aldosterone and corticosterone plus GR inhibition further indicate that MR nuclear localisation is not sufficient to determine a uniform biological output. Both conditions increased nuclear MR signal in cardiac fibroblasts, excluding insufficient receptor nuclear engagement as a simple explanation for their divergent effects. However, ligand-dependent MR conformation, differential co-regulator recruitment, distinct chromatin occupancy or altered functional interactions between MR and antagonised GR could generate different biological outputs. Direct characterisation of these aspects may explain how cardiomyocyte-proliferative MR signalling can be separated from adverse stromal programmes.

Importantly, the effects of corticosteroid receptor redirection were also detectable in adult myocardial tissue. Myocardial infarction increased circulating corticosterone levels, and genetic or pharmacological inhibition of GR enhanced cardiomyocyte MR nuclear localisation in the injured adult heart. In uninjured adult murine myocardial slices, finerenone abolished the increase in cardiomyocyte cell-cycle activity induced by corticosterone plus GR antagonism, demonstrating that the redirected response remains MR-dependent in mature myocardium. Corticosterone plus GR antagonism also enhanced cardiomyocyte cell-cycle activity following injury and elicited analogous responses in porcine and human myocardial slices. Living myocardial slices preserve tissue architecture, multicellular composition and extracellular matrix organization, providing an *ex vivo* platform in which corticosteroid responses can be examined in mature myocardium (Watson et al. 2019; Caliandro et al. 2025; Husetić et al. 2026). Together, these findings indicate that the functional GR–MR hierarchy identified in neonatal murine cardiomyocytes persists in adult myocardial tissue, while the analogous response observed in porcine and human myocardium supports its translational relevance.

From a therapeutic perspective, these findings suggest that modulating receptor hierarchy may offer a more selective strategy than simply increasing or decreasing hormone availability. Systemic activation of MR by aldosterone is unlikely to represent a viable regenerative strategy because of its established profibrotic, proinflammatory, hypertrophic and arrhythmogenic effects (Gravez et al. 2013; Richardson et al. 2016; Ayuzawa and Fujita 2021). Conversely, broad MR antagonism may suppress both pathological stromal programmes and potentially favourable cardiomyocyte responses. Our findings provide proof of principle that GR antagonism can redirect endogenous glucocorticoid signalling towards an MR-dependent cardiomyocyte-proliferative output in both neonatal cardiomyocytes and adult myocardial tissue. In multicellular cultures, this response occurs without reproducing the fibroblast activation induced by aldosterone. We thus suggest receptor hierarchy as a potentially tractable level of therapeutic control. Spatially and temporally restricted approaches, including cardiomyocyte-targeted delivery (Wang et al. 2018; Hajipour et al. 2019; Saludas et al. 2021; Smith and Edelman 2023), may ultimately help capture the favourable cardiac output of this pathway while limiting systemic and stromal consequences.

In conclusion, our findings identify functional GR-MR hierarchy and cellular context as key determinants of corticosteroid decoding in the heart. The same endogenous ligand can favour cardiomyocyte cell-cycle exit or proliferative competence depending on GR availability and on the cellular compartment in which MR is engaged. This work therefore reframes endogenous corticosteroids as context-dependent endocrine signals and provides proof of principle that their cardiac output can be redirected towards cardiomyocyte proliferation without necessarily inducing fibroblast activation.

## MATERIALS AND METHODS

### Animal experiments

All animal procedures were performed in accordance with institutional guidelines of the University of Bologna (Bologna, Italy), Cogentech/IFOM (Milan, Italy), University Medical Center (Amsterdam, the Netherlands) and Weizmann Institute of Science (Rehovot, Israel), and complied with applicable national and international regulations.

Mice were maintained on a C57BL/6 background under controlled environmental conditions, with temperature maintained at 20-25 °C, relative humidity at 40-60% and a 12-hours light/dark cycle. Cardiomyocyte-specific GR knockout mice (GR-cKO) were generated by crossing mice expressing Cre recombinase under the control of the Myh6 promoter with mice carrying loxP sites flanking exons 3 and 4 of the GR gene, as previously described (Pianca et al. 2022; Da Pra et al. 2026). GR^flox/flox^ littermates were used as controls. Genotyping primers are listed in **Supplementary Table 1**. Unless otherwise specified, data from male and female mice were pooled.

### Human studies

Approval for studies on human tissue samples was obtained from the Medical Ethics Committee of the University Medical Center, Amsterdam, the Netherlands (METC-2024.0643).

### Neonatal and juvenile cardiomyocyte isolation and culture

Primary neonatal cardiac cells were isolated from postnatal day 0-1 (P0-P1) mouse hearts by enzymatic digestion with pancreatin (P1750, Sigma-Aldrich) and collagenase A (10103586001, Roche), as previously described (Bongiovanni et al. 2024). Juvenile cardiomyocytes were isolated from postnatal day 7 (P7) hearts by anterograde perfusion with digestion enzymes containing collagenase A (10103586001, Roche), trypsin (T8003, Sigma-Aldrich) and protease (P5147, Sigma-Aldrich), as previously described (Da Pra et al. 2026).

Cells were plated on 0.1% gelatin-coated culture wells (G1393, Sigma-Aldrich) and maintained in DMEM/F12 medium (AU-L0092, Aurogene) supplemented with L-glutamine (G7513, Sigma-Aldrich), sodium pyruvate (11360070, Gibco), MEM non-essential amino acids (11140050, Gibco), penicillin/streptomycin (ECB3001D, Euroclone), 5% horse serum (16050122, Gibco) and 10% fetal bovine serum (10270106, Gibco), hereafter referred to as complete medium. Cells were cultured at 37 °C and 5% CO₂ and allowed to adhere for 48 h.

After adhesion, complete medium was replaced with FBS-deprived complete medium containing the indicated treatments: corticosterone (27840, Sigma-Aldrich), dexamethasone (D1756, Sigma-Aldrich), finerenone (HY-111372, MedChemExpress), mifepristone/RU486 (M8046, Sigma-Aldrich) or aldosterone (A9477, Sigma-Aldrich), at the concentrations specified in the figure legends. Treatments were performed for 48 hours unless otherwise indicated. For BrdU incorporation assays, BrdU (10 μM; B5002, Sigma-Aldrich) was added together with the treatments.

### Cardiomyocyte and stromal cell enrichment

To obtain cardiomyocyte-enriched and stromal-enriched cultures from P1 and P7 hearts, cardiac cell suspensions were separated using the MACS Neonatal Heart Dissociation/Isolation System (130-100-825, Miltenyi Biotec), according to the manufacturer’s instructions. Cardiomyocyte-enriched and stromal-enriched fractions were centrifuged, resuspended in culture medium and seeded for subsequent experiments.

### Time-lapse imaging

Neonatal cardiomyocyte division and binucleation were assessed by time-lapse imaging, as previously described (Pianca et al. 2022; Da Pra et al. 2026). Briefly, 48 hours after isolation, P1 GR-cKO cardiac cultures were incubated for 20 minutes with 10 nM tetramethylrhodamine ethyl ester (TMRE; 87917, Sigma-Aldrich), a mitochondrial fluorescent dye used to visualize cardiomyocytes. Cells were then treated with corticosterone (10 nM; 27840, Sigma-Aldrich) in FBS-deprived complete medium.

Live-cell imaging was performed using a Nikon Eclipse Ti2 widefield fluorescence microscope. Images were acquired at ×20 or ×40 magnification every 20 min for 24 h. Cardiomyocyte division was defined as karyokinesis followed by cytokinesis, whereas binucleation was defined as karyokinesis without completion of cytokinesis.

### *In vivo* corticosterone administration

To assess the effects of corticosterone during early postnatal development, water-soluble corticosterone-HBC complex (C174, Sigma-Aldrich) was administered at 100 μg/ml in the drinking water of lactating mothers of GR-cKO and control pups, as previously described (Da Pra et al. 2026). This administration route was selected to minimize handling-related stress. Fresh corticosterone-containing solutions were prepared daily and provided ad libitum from postnatal day 0-1 (P0-P1) to postnatal day 7 (P7).

### Myocardial infarction

Myocardial infarction was induced in adult 3-month-old mice by permanent ligation of the left anterior descending coronary artery. Mice were anaesthetized with isoflurane (Abbott Laboratories), intubated and mechanically ventilated. A left lateral thoracotomy was performed at the fourth intercostal space, and the intercostal muscles were bluntly dissected to expose the heart. The left anterior descending coronary artery was ligated, the thoracic wall was closed with 6.0 non-absorbable silk sutures and the skin incision was sealed with tissue adhesive. Mice were maintained on a warming pad until recovery.

Cardiac functional impairment was verified 1 or 2 days after myocardial infarction, and mice with comparable functional decline were included for subsequent analyses at the time points indicated in the main text and figure legends.

### Serum corticosterone measurement

Serum corticosterone levels were measured 24 hours after myocardial infarction in adult control and GR-cKO mice. Blood samples were collected by decapitation and centrifuged at 2000g for 10 minutes to isolate serum. Circulating corticosterone concentrations were quantified using a corticosterone ELISA kit (ADI-900-097, Enzo Life Sciences), according to the manufacturer’s instructions.

Absorbance was measured at 405 nm using a Tecan Spark multimode microplate reader. Corticosterone concentrations were calculated from a standard curve with a detection range of 32-20000 pg/ml.

### Preparation and culture of living myocardial slices

Human, swine and murine left ventricular myocardial tissue was sliced using a vibratome (Leica VT1200S or Campden Instruments 7000SMZ), as previously described (Watson et al. 2017). Briefly, myocardial tissue blocks were sectioned in cold cardioplegic solution. Slices were then allowed to recover in recovery solution at room temperature before being transferred to air-liquid interface culture.

Myocardial slices were cultured in M199-based medium (M4530, Sigma-Aldrich) supplemented with 1.5% penicillin-streptomycin (15070063, Gibco) and 1% insulin-transferrin-selenium (41400045, Gibco). Medium was replaced every 24 hours. Slices were treated with corticosterone (CORT; 27840, Sigma-Aldrich), RU486/mifepristone (M8046, Sigma-Aldrich), finerenone (HY-111372, MedChemExpress), or the indicated combinations, at the concentrations indicated in the figure legends.

To induce *ex vivo* myocardial injury, murine myocardial slices were subjected to cryoinjury. A copper filament was immersed in liquid nitrogen and then gently applied to the tissue surface for 2-3 seconds to generate a localized injury, as previously described (Abbas et al. 2024). Slices were subsequently cultured as described above.

### Immunofluorescence staining of cultured cells and tissue sections

Cultured cells were fixed with 4% paraformaldehyde in PBS for 20 minutes at room temperature. Paraffin-embedded heart sections were deparaffinized in xylene and rehydrated through decreasing ethanol concentrations. Heat-induced antigen retrieval was performed in 10 mM Tris-HCl, 1 mM EDTA and 0.05% Tween-20, followed by gradual cooling.

Cultured cells and tissue sections were permeabilized with 0.5% Triton X-100 in PBS for 5 minutes at room temperature. For BrdU staining, DNA hydrolysis was performed using 2 M HCl for 30 minutes at 37 °C. Non-specific antibody binding was blocked for 1 hour at room temperature using PBS containing 5% BSA (A9418, Sigma-Aldrich) and 0.1% Triton X-100 (X100, Sigma-Aldrich).

Samples were incubated overnight at 4 °C with primary antibodies diluted in PBS containing 3% BSA and 0.1% Triton X-100. The following primary antibodies were used: anti-cardiac troponin T (cTnT; 1:500, ab33589, Abcam), anti-cardiac troponin I (cTnI; 1:500, ab47003, Abcam), anti-Ki67 (1:75, ab16667, Abcam), anti-BrdU (1:40, G3G4, DSHB), anti-Aurora B kinase (1:50, 611082, BD Transduction Laboratories), anti-MR (1:50, MABS496, Sigma-Aldrich), anti-vimentin (1:100, ab92547, Abcam) and anti-αSMA (1:500, BK19245T, Cell Signalling).

After primary antibody incubation, samples were washed and incubated for 1 hour at room temperature with the following fluorescent secondary antibodies diluted 1:200 in PBS containing 1% BSA and 0.1% Triton X-100: anti-mouse Alexa Fluor 488 (115-545-003, Jackson ImmunoResearch), anti-rabbit Alexa Fluor 488 (111-545-003, Jackson ImmunoResearch), anti-mouse Cy3 (115-165-003, Jackson ImmunoResearch) and anti-rabbit Cy3 (111-165-003, Jackson ImmunoResearch).

Nuclei were counterstained with DAPI (D9542, Sigma-Aldrich) for 10 minutes at room temperature. Cultured cells were imaged using a Nikon Eclipse Ti2 widefield fluorescence microscope and analysed with NIS-Elements 5.21 software (Nikon). Tissue sections were mounted with DPX mountant (06522, Sigma-Aldrich), coverslipped, sealed and imaged using an Olympus VS200 slide scanner.

### Assessment of cardiomyocyte cross-sectional area

Cardiomyocyte cross-sectional area was measured in P7 heart sections using wheat germ agglutinin (WGA) staining. Sections were deparaffinized, rehydrated and washed in PBS before incubation with Alexa Fluor 488-conjugated WGA (1:200, 29022-1, Biotium) for 45 minutes. Sections were washed three times in PBS, mounted with aqueous mounting medium (Mount Quick Aqueous, 2435489, Bio-Optica) and imaged using a Nikon Eclipse Ts2 microscope.

Cross-sectional area was quantified using ImageJ software (National Institutes of Health). Only transversely oriented cardiomyocytes were included in the analysis.

### Estimation of cardiomyocyte number in juvenile hearts

Cardiomyocyte number in P7 hearts was estimated by stereological analysis. Heart weight was multiplied by the density of muscle tissue (1.06 g/ml) to estimate total heart volume. The total heart volume was then multiplied by the fraction occupied by cardiomyocytes, calculated from cTnT-stained heart sections, to estimate total cardiomyocyte volume per heart.

Average cardiomyocyte length was determined from isolated P7 cardiomyocytes using ImageJ, as previously described (Pianca et al. 2022). Average cardiomyocyte volume was calculated by multiplying cardiomyocyte cross-sectional area by average cardiomyocyte length. The total number of cardiomyocytes per heart was estimated by dividing the total cardiomyocyte volume by the average cardiomyocyte volume.

### Transient gene knockdown

For transient gene silencing, SMARTpool siRNAs targeting mineralocorticoid receptor (MR/Nr3c2), androgen receptor (AR/Nr3c4), oestrogen receptor-α (ERα/Esr1), oestrogen receptor-β (ERβ/Esr2), G protein-coupled oestrogen receptor 1 (GPER1/Gper1) and progesterone receptor (PR/Pgr) were delivered to P1 cardiac cultures 24 hours after seeding using Lipofectamine 2000 (11668-019, Thermo Fisher Scientific), according to the manufacturer’s instructions. Lipofectamine 2000 was used at the lowest recommended concentration.

Gene knockdown was assessed 48 hours after transfection by gene-expression analysis. For proliferation assays, medium was replaced after transfection with FBS-deprived complete medium containing corticosterone for 48 hours. A pool of scrambled siRNAs was used as negative transfection control in all experiments.

### Gene-expression analysis

Total RNA was extracted using the NucleoSpin RNA II kit (Macherey-Nagel), according to the manufacturer’s instructions. RNA concentration and purity were assessed using a NanoDrop spectrophotometer (N1000, Thermo Fisher Scientific). RNA was reverse transcribed into cDNA using the RevertAid RT kit (K1691, Thermo Fisher Scientific), according to the manufacturer’s protocol.

Real-time quantitative PCR was performed using Fast SYBR Green PCR Master Mix (4309155, Applied Biosystems) on a QuantStudio 5 Flex real-time PCR system (Applied Biosystems). Relative gene expression was calculated using the ΔΔCt method and normalized to HPRT1. Oligonucleotide sequences used for quantitative PCR are listed in **Supplementary Table 2**.

### Quantification and statistical analysis

Statistical analyses were performed using GraphPad Prism. Data are presented as mean ± s.e.m. or median, as indicated in the figure legends. When data distribution and experimental design allowed parametric testing, comparisons between two groups were performed using two-sided Student’s t-test, whereas comparisons among multiple groups were performed using one-way ANOVA followed by Šidák’s or Tukey’s multiple-comparisons test, as specified in the figure legends.

A P value < 0.05 was considered statistically significant. Statistical significance is indicated in the figures as follows: *P < 0.05, **P < 0.01, ***P < 0.001 and ****P < 0.0001.

## ACKNOWLEDGMENTS

We acknowledge financial support under the National Recovery and Resilience Plan (NRRP), Mission 4, Component 2, Investment 1.1, Call for tender No. 104 published on 2.2.2022 by the Italian Ministry of University and Research (MUR), funded by the European Union - NextGenerationEU- Project Title “Fine-tuning corticosteroid signalling as a novel therapeutic strategy for heart regeneration” - CUP J53D23012310006 - Grant Assignment Decree No. 1065 adopted on 18/07/2023 by the Italian Ministry of University and Research (MUR).

This work was also supported by the Italian Ministry of Health (grant no. RC24000858-2795994 ex 2790614 to G.D’U.).

## AUTHOR CONTRIBUTIONS

F.S. and G.D’U. designed the experiments. F.S. carried out most of the experiments and analysed the data. S.B. and G.D’U. performed *in vivo* murine procedures. F.S., R.C. and A.H. performed living myocardial slices experiments. S.B., S.D.P., I.D.B., C.B., C.M., N.P., S.M., A.C. performed immunofluorescences and gene expression analyses. R.T. helped with immunofluorescence image acquisition and time-lapse imaging. M.L., C.V., and M.M.G. supervised the experiments done by their laboratory members, and G.D’U. supervised the entire project. F.S. and G.D’U. wrote the manuscript, with editing contributions from all authors.

