## Supplementary Figures for "Hierarchical and context-dependent GR-MR signalling governs endogenous corticosteroid decoding in the heart"

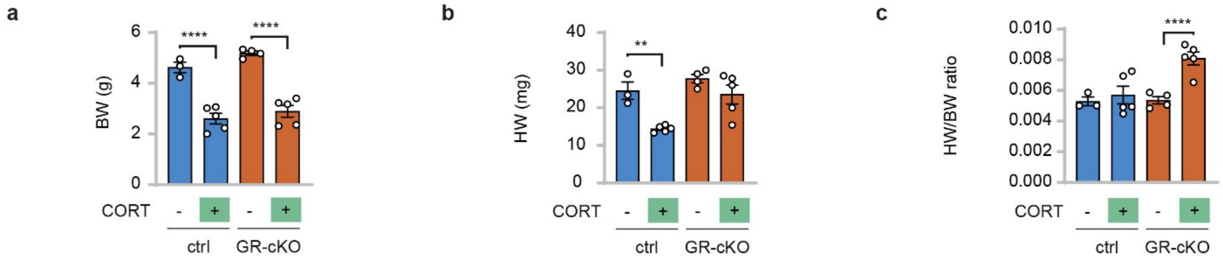


**Supplementary Figure 1. Postnatal growth and heart size after *in vivo* corticosterone administration.** (**a-c**) Water-soluble corticosterone–HBC complex (100 µg/ml) was administered in the drinking water of lactating mothers during the first week of pups postnatal life. Body weight (BW, **a**), heart weight (HW, **b**) and heart-weight-to-body-weight ratio (HW/BW, **c**) of pups were measured at the end of treatment at postnatal day 7 (P7) (n = 17 mice). Data are presented as mean ± s.e.m. Statistical significance was assessed by one-way ANOVA followed by Šidák's multiple-comparisons test. *P < 0.05, **P < 0.01, ***P < 0.001, ****P < 0.0001.

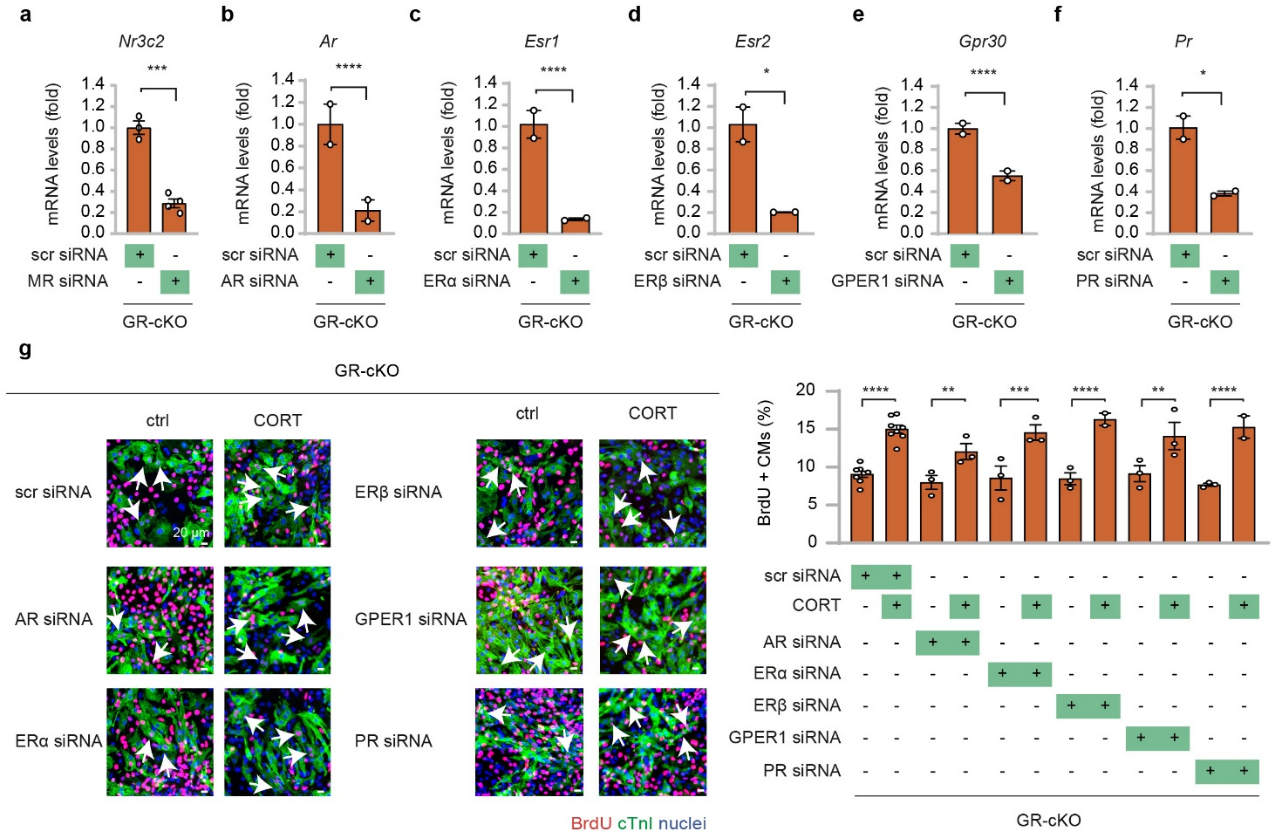


**Supplementary Figure 2. Steroid hormone receptor silencing in GR-deficient neonatal cardiomyocyte cultures.** (**a**–**f**) Validation of siRNA-mediated silencing of steroid hormone receptors in neonatal primary cardiac cultures isolated from cardiomyocyte-specific GR knockout (GR-cKO) mice. Expression of genes coding for mineralocorticoid receptor (*Nr3c2*; **a**), androgen receptor (*Ar*; **b**), oestrogen receptor-α (*Esr1*; **c**), oestrogen receptor-β (*Esr2*; **d**), G protein-coupled oestrogen receptor 1 (*Gpr30*; **e**) and progesterone receptor (*Pr*; **f**) was assessed after transfection with receptor-specific siRNAs (MR, AR, ERα, ERβ, GPER1 and PR siRNAs, respectively). Scrambled siRNA (scr) was used as transfection control (n = 7 samples for *Nr3c2*; n = 4 samples for *Ar*, *Esr1*, *Esr2*, *Gpr30* and *Pr*); (**g**) AR, ERα, ERβ, GPER1 and PR were individually silenced by siRNA transfection in GR-cKO neonatal primary cardiac cultures. Cardiomyocyte proliferation was assessed by BrdU incorporation after corticosterone (CORT, 10 nM) treatment (n = 42 samples; 11427 CMs analysed). Representative images are shown; arrows indicate proliferating CMs. Scale bars, 20 µm. Data are presented as mean ± s.e.m. Statistical significance was assessed by two-sided Student's t-test in **a**–**f**, and by one-way ANOVA followed by Šidák's multiple-comparisons test in **g**. *$P < 0.05$, **$P < 0.01$, ***$P < 0.001$, ****$P < 0.0001$.

### Supplementary Videos

**Supplementary Video 1. Time-lapse imaging of cell division in neonatal cardiomyocytes.** Representative time-lapse movie showing a GR-cKO neonatal cardiomyocyte undergoing karyokinesis followed by cytokinesis after corticosterone treatment. Primary cardiac cells were isolated from P1 GR-cKO mice, cultured in vitro for 48 hours, treated with corticosterone, labelled with TMRE (green) to identify cardiomyocytes, as described in the Methods, and imaged for 24 hours at 20-minutes intervals.

**Supplementary Video 2. Time-lapse imaging of binucleation in neonatal cardiomyocytes.** Representative time-lapse movie showing a P1 GR-cKO neonatal cardiomyocyte undergoing karyokinesis without cytokinesis, resulting in binucleation, after corticosterone treatment. Primary cardiac cells were isolated from P1 GR-cKO mice, cultured in vitro for 48 hours, treated with corticosterone, labelled with TMRE (green) to identify cardiomyocytes, as described in the Methods, and imaged for 24 hours at 20-minutes intervals.

### Supplementary Tables

**Supplementary Table 1.** Sequences of the primers used for mouse genotyping analyses.

| Gene | Forward primer | Reverse primer |
|---|---|---|
| $GR^{flox}$ | ATGCCTGCTAGGCAAATGAT | TTCCAGGGCTATAGGAAGCA |
| αMyHC-Cre | ATGACAGACAGATCCCTCCTATCTCC | CTCATCACTCGTTGCATCATCGAC |
| αMyHC-Cre (internal control) | CAAATGTTGCTTGTCTGGTG | GTCAGTCGAGTGCACAGTTT |

**Supplementary Table 2.** Sequences of the primers used to analyse mRNA levels by real-time (rt)PCR.

| Gene | Forward primer | Reverse primer |
|---|---|---|
| *Hprt1* | ATAAGCCAGACTTTGTTGG | ATAGGACTCCAGATGTTTCC |
| *Nr3c2* (MR) | TGCATGATTTGGTGAATGAC | CTTCCTGTGAAAGTAAAGGG |
| *Nr3c4* (AR) | TCTTCAGCATTATTCCAGTG | GATCGAGTTCCTTGATGTAG |
| *Esr1* (ERα) | CAAGGTAAATGTGTGGAAG | GTGTACACTCCGGAATTAAG |
| *Esr2* (ERβ) | CTCAACTCCAGTATGTACCC | CATGAGAAAGAAGCATCAG |
| *Gpr30* (GPER1) | GATCTAGGGAGAAAGCCAT | CCTGTTAGTCTCAGAAAACC |
| *Pgr* (PR) | AACTCACAAAACTTCTCGAC | ACTTTTTGTGAAAGAGGAGC |
